# A cell fractionation and quantitative proteomics pipeline to enable functional analyses of cotton fiber development

**DOI:** 10.1101/2024.08.08.603566

**Authors:** Youngwoo Lee, Heena Rani, Eileen L. Mallery, Daniel B Szymanski

## Abstract

Cotton fibers are aerial trichoblasts that employ a highly polarized diffuse growth mechanism to emerge from the developing ovule epidermis. After executing a complicated morphogenetic program, the cells reach lengths over 2 cm and serve as the foundation of a multi-billion-dollar textile industry. Important traits such as fiber diameter, length, and strength are defined by the growth patterns and cell wall properties of individual cells. At present, the ability to engineer fiber traits is limited by our lack of understanding regarding the primary controls governing the rate, duration, and patterns of cell growth. To gain insights into the compartmentalized functions of proteins in cotton fiber cells, we developed a label-free liquid chromatography mass spectrometry method for systems level analyses of fiber proteome. Purified fibers from a single locule were used to fractionate the fiber proteome into apoplast (APO_T_), membrane-associated (p200), and crude cytosolic (s200) fractions. Subsequently, proteins were identified, and their localizations and potential functions were analyzed using combinations of size exclusion chromatography, statistical and bioinformatic analyses. This method had good coverage of the p200 and apoplast fractions, the latter of which was dominated by proteins associated with particulate membrane-enclosed compartments. The apoplastic proteome was diverse, the proteins were not degraded, and some displayed distinct multimerization states compared to their cytosolic pool. This quantitative proteomic pipeline can be used to improve coverage and functional analyses of the cotton fiber proteome as a function of developmental time or differing genotypes.

## Introduction

*Gossypium hirsutum* (*G. hirsutum*) is the most widely grown cotton, ranking among the world’s most economically important crop species, and provides the largest source of renewable textiles (over $6 billion in raw product, FAOSTAT, http://www.fao.org). This global textile economy is based solely on the growth and morphogenesis of individual fiber cells that emerge from the developing seed coat. At about 1 day before flower opening and anthesis, conserved transcriptional control pathways drive subsets of epidermal precursors into the trichoblast fate (Szymanski et al., 2000; Wang et al., 2020b). Subsequently the individual cells execute a complicated and poorly understood morphogenetic program (Delmer, 1999; Haigler et al., 2012; Kim and Triplett, 2001; Qin and Zhu, 2011) that includes an initial tapering phase that reduces fiber diameter (Applequist et al., 2001; Graham and Haigler, 2021; Yanagisawa et al., 2022).

Daily phenotyping of fiber traits and cell wall molecular features reveal a progressively slowing fiber elongation rate during early stages of development (Wilson, 2024). Well after fiber elongation slows, fibers execute a multi-phased transition to the synthesis of a cellulose-rich secondary cell wall that eventually consumes much of the cell cytoplasm prior to cell death, desiccation, and the opening of the mature boll. Domestication and breeding have generated modern cultivars with narrow fibers, extended developmental phases of cell elongation, and twisted morphologies that enable dried fibers to be spun into a usable product. Because the genetic diversity in the cottons is relatively low, future opportunities reside in the combined use of genetic engineering and breeding to improve fiber traits and yield. Even when dealing with individual cells this is a complicated engineering challenge.

Fiber development includes complex developmental interactions among central metabolism and turgor regulation (Ruan et al., 2001; Tuttle et al., 2015). The biomechanics of how turgor pressure interacts with the material properties of the growing trichoblast cell wall is not completely understood. Aerial trichoblasts, like most plant cells (Gu and Rasmussen, 2022), employ a diffuse growth strategy that couples persistent synthesis and assembly of a cellulose-rich wall with throughout the expanding cell surface (O’Kelley, 1952; Ryser, 1977; Yanagisawa *et al*., 2022). This enables the cell to not only maintain the wall strength needed to avoid cell rupture by tensile stress (Fujita et al., 2013), but also maintain anisotropic wall material properties that enable polarized elongation with minimal radial (Baskin, 2005; Graham and Haigler, 2021; Yanagisawa et al., 2015; Yanagisawa *et al*., 2022). A microtubule-patterned cellulose synthesis system operates throughout fiber development to promote persistent axial elongation and cell diameter control at the cell apex (Seagull, 1986; Wilson, 2024; Yanagisawa *et al*., 2022).

The microtubule-cellulose control module is just one part of a much broader network of cellular systems that dictate the rates and patterns of cell expansion. As just one example, matrix polysaccharides like the pectins, homogalacturonans, and rhamnogalacturonans are major components of the growing cell wall (Avci et al., 2013; Swaminathan S, 2024), and their material properties can have strong effects on the rates and patterns of growth in leaf trichoblasts (Yanagisawa *et al*., 2015) and cotton fibers (Wilson, 2024). Pectin biosynthesis, transport, and modification involve dozens of genes that must be tightly regulated during growth, so that the thickness and material properties of the wall are maintained during a growth phase that proceeds for weeks (Meinert and Delmer, 1977; Schubert et al., 1973). There is a similar level of complexity to mediate the transition to very different physiology of the cell as it rearranges its metabolism, endomembrane systems, and glycosyl transferase repertoire to form the secondary wall (Hoffmann et al., 2021; Zhong et al., 2019).

There are emerging opportunities to discover key molecules and systems-level interactions that govern fiber traits. Forward genetic screens are complicated by genetic redundancy in tetraploid cottons; however, *G. hirsutum.* fiber morphology mutants have enabled discovery of cytoskeleton related genes that affect fiber length and twist (Wan et al., 2016; Zang et al., 2021; Zhang et al., 2021). Because fiber cells develop in a synchronous manner and tens of thousands of fiber cells emerge from a single developing seed coat (Bowman et al., 2001), it is possible to conduct transcriptomic and metabolomic analyses using either purified fibers or developing ovules of different genotypes to predict pathways and genes that underlie fiber quality (Qin et al., 2019; Rapp et al., 2010; Tuttle *et al*., 2015; Wang et al., 2015; You et al., 2023).

Cotton fiber proteomic analyses are especially valuable because they can provide direct measurements of homoeolog utilization, protein abundance, and post-translational modifications in polyploid species (Hu et al., 2014; Soltis et al., 2016; Wang et al., 2020c). Gel-free workflows and the increased sensitivity of modern spectrometers have enabled the quantification of thousands, rather than dozens, of proteins in a typical experiment (Hu et al., 2015; Hu et al., 2013; Lee and Szymanski, 2021; Yang et al., 2023; Yang et al., 2008). Proteins with an altered abundance in response to domestication (Bao et al., 2011; Hu *et al*., 2013; Qin et al., 2017), hormone or inhibitor responses (Wang et al., 2020a; Yang *et al*., 2023), and developmental stages (Jiang et al., 2022; Mujahid et al., 2016; Zhou et al., 2019) have been identified. There are also growing opportunities to use quantitative proteomics and correlation profiling as a systems-level phenotyping tool that analyzes multimerization state, localization, and protein complex composition (Lee and Szymanski, 2021; McBride et al., 2019).

The primary focus of this paper is to develop a robust quantitative proteomics pipeline using purified fibers isolated from a single locule in the developing boll to enable deeper proteomic phenotyping. The workflow combines cell fractionation with label-free shotgun proteomics to increase protein coverage and generate crude estimates of protein localization. Because of the central importance of cell wall remodeling (Avci *et al*., 2013; Pettolino et al., 2022; Yanagisawa *et al*., 2022) and developmentally regulated apoplastic sugar metabolism (Ruan *et al*., 2001) during fiber development we included a gentle locule wash step to target loosely bound extracellular proteins. Similar strategies to analyze the apoplast in other plant systems detects both soluble secreted proteins and those associated with lipid-enclosed extracellular vesicles. Here we report on a diverse class of proteins in the apoplastic fraction. Soluble apoplastic proteins are not proteolytic remnants of cytosolic proteins, and many display unique multimerization states in the apoplast compared with the intracellular pool. Most proteins in the apoplastic proteins are associated with lipid-enclosed vesicles in the apoplastic fraction.

The apoplastic vesicle-associated proteome closely resembles that of crude microsomes; however, there are hundreds of proteins and protein complexes that are significantly enriched in the apoplastic fraction. We provide an example database in which well-annotated protein data are searchable as a function of subcellular location and relative abundance, and a method that can be adapted to analyze developmental progressions of differences among genotypes.

## Results

### Cell fractionation and quantitative proteomics of purified fibers

We sought to create a method that would enable the quantitative analysis of protein abundance and localization in developing cotton fibers dissected from individual locules from a developing boll (Figure 1A). Almost all fibers on the isolated locules at 9 day post anthesis (DPA) were intact (Figure 1B) and showed vesicle-like particles on the surface of fiber (Figure 1B (4) and (5)). To increase protein coverage and analyze protein localization in a crude manner, a total apoplast fraction (APO_T_) of loosely bound proteins was obtained after careful dissection and dunking of a single locule, then crude microsomal proteins in the high-speed pellet (p200), and total soluble (s200) fraction enriched in soluble cytosolic proteins were prepared. Due to fragility of the developing fibers, we generated the APO_T_ fraction of loosely bound apoplastic proteins by gently transferring the dissected locule to a microsome isolation buffer (MIB) solution with gentle rocking for 10 minutes. When the locules were analyzed by confocal microscopy after staining with propidium iodide to detect broken cells, the samples before (water) or after the 10 min incubation procedure were indistinguishable with little evidence for broken cells (Figure 1C), and the fraction was not dominated by broken cells. After the APO_T_, fibers were manually dissected from the seeds, homogenized, filtered, low-speed spun, and ultracentrifuged to generate a total soluble protein (s200) and a crude microsomal fraction (p200). The centrifugation protocol does not sediment large protein complexes like Rubisco (Aryal et al., 2014) and this protocol will generate a washed crude microsome fraction that is highly enriched in true-membrane associated proteins (McBride et al., 2017).

**Figure 1.**
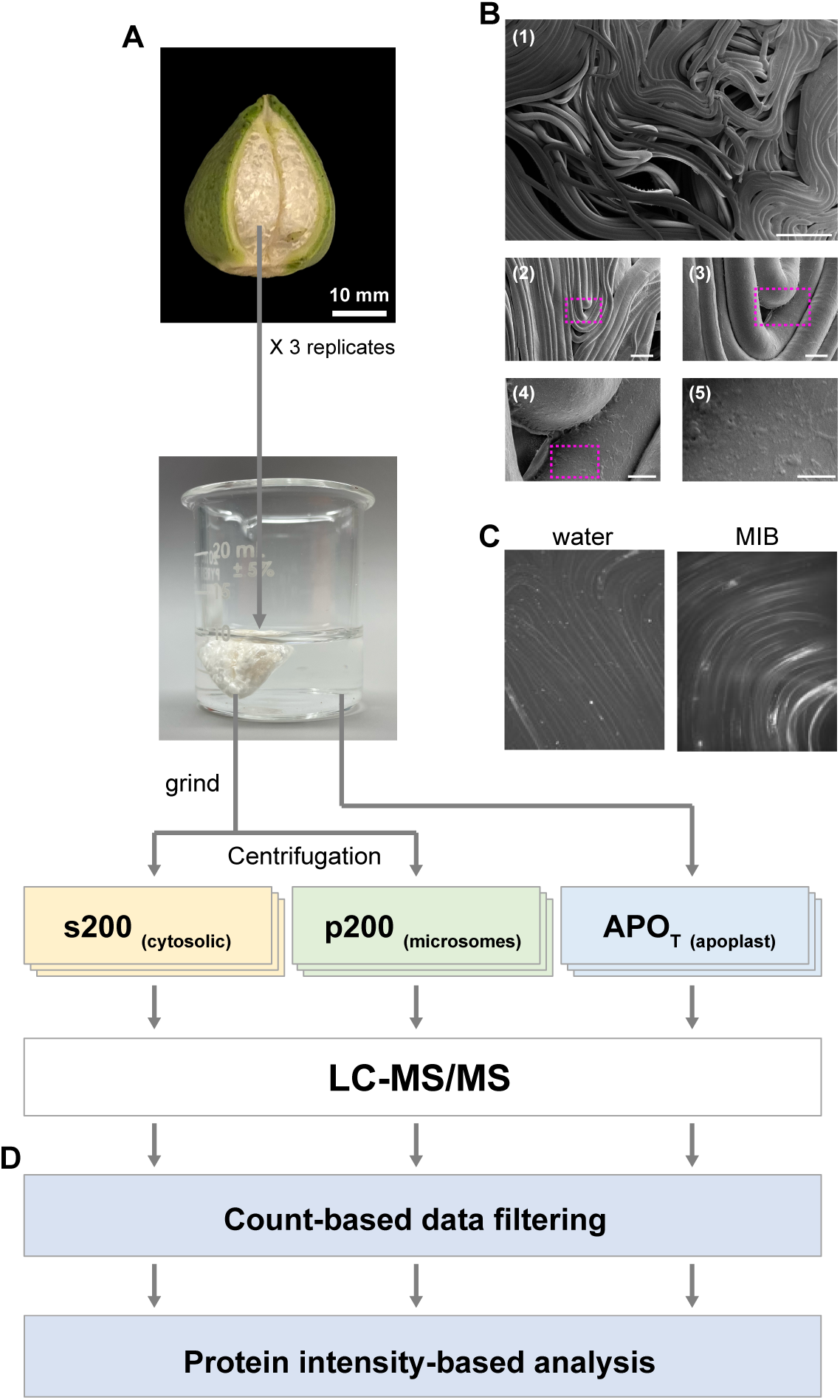
Discovery of apoplast-enriched cotton fiber proteins via cell fractionation and quantitative mass spectrometry. **A,** Protein isolation method from each locule at 9 DPA boll for shotgun proteomics. **B,** Scanning electron microscopy analysis of developing cotton fibers at 15 DPA. (1) Scanning electron micrograph demonstrating intact fibers. Bar, 200 μm. (2) to (5) Zoom-in views of fiber surface. Bar, 50 μm (2), 10 μm (3), 5 μm (4), and 2 μm (5). Boxed regions with pink dotted lines were zoomed in. **C,** Propidium iodide-stained images of the intact locules before (Water) and after (MIB) 10 minute-dipping in MIB via 20X confocal microscope. **D,** A focus on proteins with a differential accumulation in the APO_T_ (apoplast). Spectral count and precursor peptide ion intensity-based quantification to predict proteins with an increased abundance in the apoplast.

The single locule protein yields of APO_T_ and p200 were sufficient for shotgun proteomics experiments (Figure 2A). To promote consistent locus ID assignments between annotated *Gossypium hirsutum* genome annotations, a locus ID conversion table is provided in Supplemental Dataset S1. Triplicate samples were processed for label free shotgun proteomics using previously established methods (Cox et al., 2014; Rudolph and Cox, 2019; Tyanova et al., 2016a; Tyanova et al., 2016b) and the raw LC-MS/MS search results (Supplemental Dataset S1) and raw LC/MS files are available at PRIDE (accession code: PXD051721). To facilitate consistent protein identifications and quantitative comparisons across all samples, the 9 LC/MS raw files from the cell fractions were searched simultaneously using MaxQuant and LFQ normalization to standardize protein intensities across samples (Cox *et al*., 2014; Tyanova *et al*., 2016a). However, s200 yield was low (Supplemental Dataset S1A) as was protein coverage (Figure 2A).

**Figure 2.**
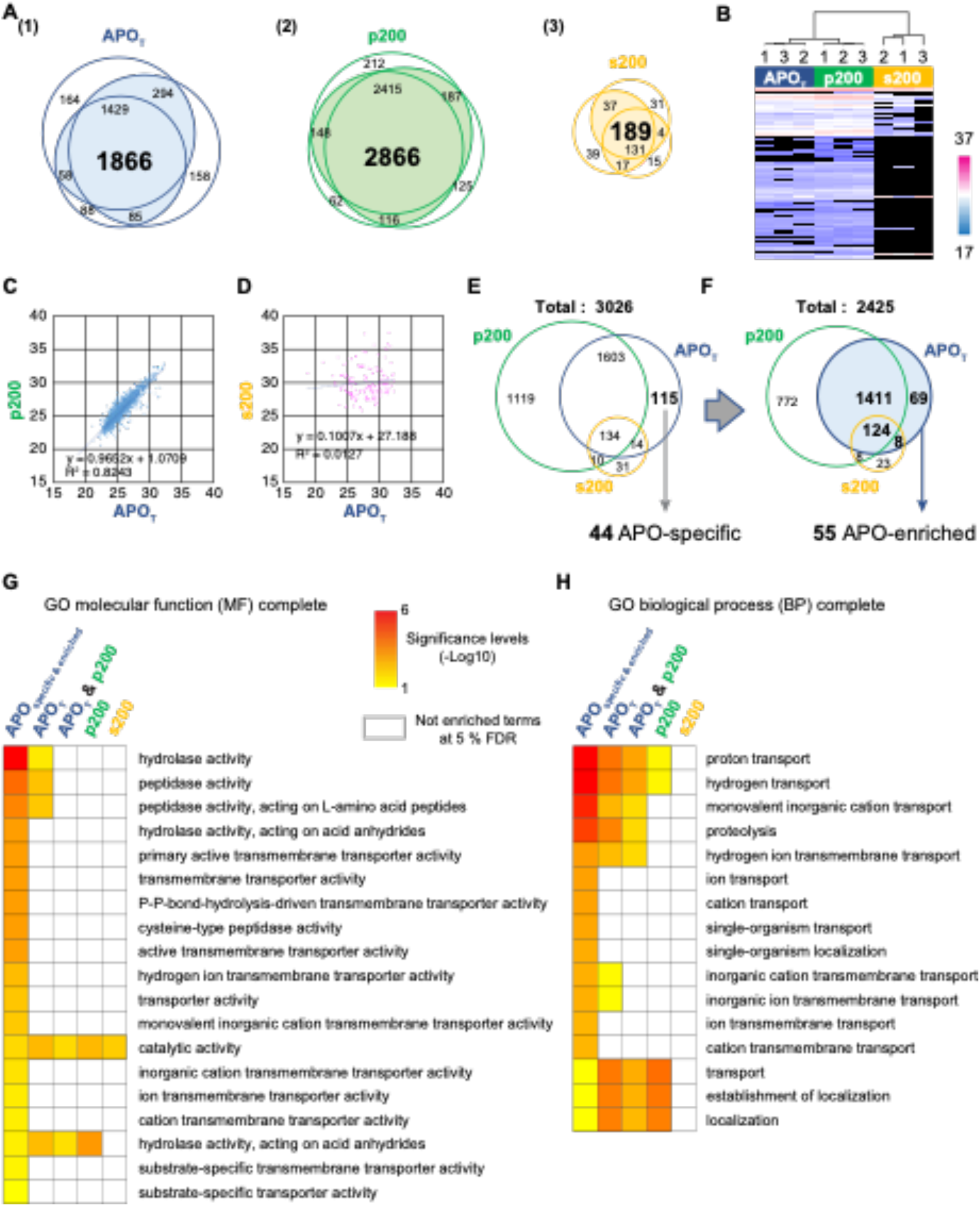
Proteome coverage, reproducibility, and compositional differences among the three cell fractions. **A,** Number of proteins reproducibly identified by MS/MS counts in the APO_TOTAL_ (1), p200 (2), and s200 (3) cellular fractions. Numbers in bold indicate reproducible proteins identified 2 out of 3 replicates. **B,** Hierarchical clustering and heatmap showing relative protein expression (log2-transformed LFQ protein intensities) of the differentially expressed proteins in the cellular fractions. Columns correspond to the three replicates of the fractions. Proteins that were not present in the datasets were rendered with black color. **C and D,** Abundance analyses of dually localized APO_T_ proteins in p200 (C) and s200 (D). log2-transformed LFQ protein intensities. **E,** Identification of apoplast-specific (APO_specific_) proteins based on MS/MS count data across the replicates and cell fraction. **F,** Protein-intensity-based identification of apoplast-enriched (APO_enriched_) proteins compared to p200 and s200 fractions. Expression levels of the quantified APO_T_ proteins (the circle with blue background color) were compared to either p200 and s200 using two-sample *t*-tests at 5 % FDR and 2-fold changes. **G and H**. GO enrichment analyses of the three cellular fractions. The agriGO v2.0 Singular Enrichment Analysis tool (http://systemsbiology.cau.edu.cn/agriGOv2/index.php) was implicated at 5% FDR. The heat maps show molecular function (G) and biological process (H) categories of the over-represented GO terms in the APO_specific_ and APO _enriched_ protein list. Significance levels of the over-represented GO terms in APO_T_, p200, and s200 protein lists, and common protein list between APO_T_ and p200 were shown in the heat maps. GO terms that were not over-represented in each dataset were boxed with white background.

3026 proteins were classified as present based on at least one unique peptide detected in two of the three replicates for a single cell fraction. The protein identifications and quantification data are provided in the Supplemental Dataset S2. The number of proteins identified within each cell fraction were reproducible (Figure 2A and 2B). The APO_T_ and p200 had similar protein compositions, and s200 was most different due in part to lower coverage (Figure 2E and 2F; Supplemental Dataset S2). The s200 sample contained only 79 enzymes, and 45 of them scattered throughout central metabolism and oxidative stress pathways (Supplemental Dataset S2E). Improved methods are needed for better coverage of this compartment. The APO_T_ and p200 fractions contained thousands of proteins including enzymes, receptor kinases, transporters, cytoskeletal proteins, kinases, transcription factors. In about 40 % of the cases the peptide data were sufficient to resolve homoeologs, while about 46 % of ambiguously assigned to both homoeologs (Supplemental Dataset S2A). Also, the quantitative data for each protein is provided both for spectral counts and LFQ-normalized precursor ion intensities (Supplemental Dataset S2A). The data can be sorted to view proteins based on their ranked estimated abundance within the cell fraction. The data can also be searched by the Arabidopsis best hit locus ID or by orthologous group number based on Phytozome V12 (Goodstein et al., 2011). As an example, cellulose synthases (CESAs), Korrigan/glycosyl hydrolase 9 (GH9), cellulose synthase interactor (CSI), companion of cellulose synthase (CC), and COBRA proteins are directly involved in patterned cellulose synthesis (Polko and Kieber, 2019). In our dataset we detect cotton orthologs for 8 of these known players (Supplemental Dataset S2F), a large family of tubulins, and microtubule-associated proteins (MAPs) that affect the dynamics and organization of the cortical microtubule array (Buschmann et al., 2010). We detected members of 59 different orthologous group proteins that relate to microtubules and may affect fiber morphogenesis (Supplemental Dataset S2F). We expect this method to generate broadly useful data about the medium to high abundance proteins in the cell that execute functions that influence fiber traits.

Unexpectedly, the composition of the APO_T_ sample resembled that of the p200 sample in terms of the detected proteins (Figure 2B) and there was considerable overlap in the molecular functions and biological processes that are predicted to be enriched in each cell fraction (Supplemental Dataset S3A, S3B, and S3D). The apoplast proteome in this study also had substantial overlap in composition of apoplastic proteomes reported previously (Supplemental Dataset S4). When the cotton protein IDs were converted to their closest Arabidopsis or rice ortholog, the cotton APO_T_ contained 54% of the proteins detected in an Arabidopsis fraction enriched in extracellular vesicles (Rutter and Innes, 2017). There was more than 40% overlap with most of the apoplast proteome surveys (Basu et al., 2006; Ge et al., 2011; Kaffarnik et al., 2009; Kwon et al., 2005; Tran and Plaxton, 2008; Trentin et al., 2015), and the lowest overlap was ∼27% (Boudart et al., 2005; Cheng et al., 2009). None of the above published datasets contained a population of apoplastic proteins with narrowly defined predicted functions or localizations, and the same is true with our cotton data here (Supplemental Dataset S4A).

The APO_T_ and p200 proteomes were strongly correlated in terms of the abundances of the dual localized proteins (Figure 2C). If APO_T_ proteins were due solely to contamination one would expect a high correlation with dual localized proteins in the s200 fraction. No such relationship was found, even though the protein populations were medium to high-abundance proteins in both cell fractions (Figure 2D). These data indicate that the protein route(s) to the apoplast (which are not known) are not linked to a particular organelle or the APO_T_ samples are highly contaminated with broken cells that selectively contribute microsome-associated proteins.

To better understand the possible contribution of broken cell contamination, we first tested for an effect of the isolation buffer on protein yield. The APO_T_ sample contained proteins that were distributed between particulate (APO_p200_) and soluble (APO_s200_) fractions, with yields of ∼64 µg/boll and ∼16 µg/boll, respectively. Similar protein yields were obtained with a wide range of buffers including deionized water (Supplemental Dataset S5). Particles in the APO_p200_ fraction were membrane-enclosed based on electron and fluorescence microscopy (see below). As an additional test a completely different APO wash method was developed to avoid removal of the locules. When the lower half of the boll capsule wall was removed and the locules were dunked in deionized water (Supplemental Figure S1A), fluorescent particles were persistently generated at the time scale of hours (Supplemental Figure S1B-S1F) and seemed unlikely to arise solely from surficial organelles fragments generated by broken cells from the capsule wall. Few to no fluorescent particles were detected when the intact inner capsule wall was similarly exposed to buffers (see methods). However, in the same assay the isolated endocarp walls that remain after locule dissection was a potent source of FM4-64 positive apoplastic vesicles. It is therefore possible that some of the APO_T_ protein from dunked locules arise from surgically damaged cells from the capsule wall. Because of this ambiguity and the difficulty in proving the apoplastic vesicles are not derived from broken cells we will refer to the membrane-enclosed particles as APO_T_-vesicles. We also detected APO_T_-vesicles in our *in vitro* ovule culture system (Supplemental Figure S2). Pellets obtained by ultracentrifuge from culture fluid in this ovule culture system were positively stained with FM4-64. This supports that those APO_T_-vesicles do not arise in broken fibers, endocarp, or capsule wall.

We next tested for proteins with an increased protein abundance in the APO_T_ fraction. We defined apoplast-specific (APO_specific_) proteins as those that had spectral counts in 2 or 3 replicates in the APO_T_ fraction and zero counts in any of the replicates from the s200 and p200 fractions (Figure 2E). 44 proteins satisfied these criteria (Supplemental Dataset S6A), and only a small fraction of them are extracellular proteins based on the SUBAcon localization prediction database (Heazlewood et al., 2007). To further test for proteins with an elevated abundance in APO_T_, relative protein abundances in in the three cell fractions were statistically analyzed using LFQ normalized precursor ion intensities (Cox *et al*., 2014). In pair wise *t*-tests between APO_T_ and p200 or s200 proteins at a false discovery rate of 5 %, 55 proteins were specifically enriched in APO_T_ (Supplemental Dataset S6B). These APO-enriched proteins are unlikely to be contaminating high abundance proteins from broken cells because their ranked relative abundance was elevated compared that in the s200 or p200 fractions (Supplemental Dataset S6B). Terms of hydrolase/peptidase and a wide variety of transmembrane transporter were enriched GO terms in the APO_specific_ and APO_enriched_ proteins, compared to APO_T_, p200 and s200 (Figure 2G and 2H, Supplemental Dataset S3E). Surprisingly, RuBisCO large and small subunits are APO_specific_ and an enzyme involved in starch synthesis, Gorai.004G128700, is APO_specific_.

This did not correspond to a clear-cut enrichment with plastid proteins in the APO_T_ fraction compared to p200, because 124 and 286 plastid proteins were detected in each cell fraction, respectively (Supplemental Dataset S2B and S2C). The SUBAcon localization patterns in the data were analyzed further after the APO_T_ fraction was centrifuged to separately analyze soluble and particulate subfractions of the apoplast.

### Analysis of protein size and protein multimerization in the apoplastic and cytosolic fractions

It is possible that the apoplast is a proteolytic compartment and we are only detecting degraded or partially degraded proteins. To address this question, apparent mass (*M*_app_) of soluble apoplastic proteins (APO_s200_) was determined using a label-free quantification of SEC column fractions (Figure 3A; Supplemental Datasets S7 and S8). We have developed robust methods for large scale analyses of protein multimerization under non-denaturing conditions (Aryal *et al*., 2014; Havugimana et al., 2012; Kristensen et al., 2012; Lee et al., 2021; Lee and Szymanski, 2021; McBride *et al*., 2019; McBride *et al*., 2017; McWhite et al., 2020; Wan et al., 2015). Reliable Gaussian-fitted peaks are generated from protein intensity values from multiple column fractions with >90 % of the proteins having identical *M*_app_ measurements between replicates (Aryal *et al*., 2014; Lee *et al*., 2021; Lee and Szymanski, 2021; McBride *et al*., 2019; McBride *et al*., 2017). Of the 205 proteins detected in the SEC separations here, 142 could be fitted to a Gaussian profile and all 46 detected in both replicates had an identical peak location and *M*_app_ value and were flagged as “replicated” (Figure 3B and 3C; Supplemental Dataset S8). To leverage the high reliability of the data, *M*_app_ values those with a Gaussian peak on one replicate were retained and flagged as “single replicate”. Proteins in the APO_s200_ fraction were not degraded. Using *R*_app_, the ratio of *M*_app_ over the calculated monomeric protein mass (*M*_mono_), as a metric protein multimerization (Liu et al., 2008), 69 proteins were predicted to be monomeric, and 61 had an *R*_app_ greater than 1.6 and were predicted to assemble into some sort of complex (Figure 3D; Supplemental Dataset S8A). Only 12 proteins, including a 20.3 kDa polygalacturonase fragment had an *R*_app_ < 0.6 and potentially was not full length.

**Figure 3.**
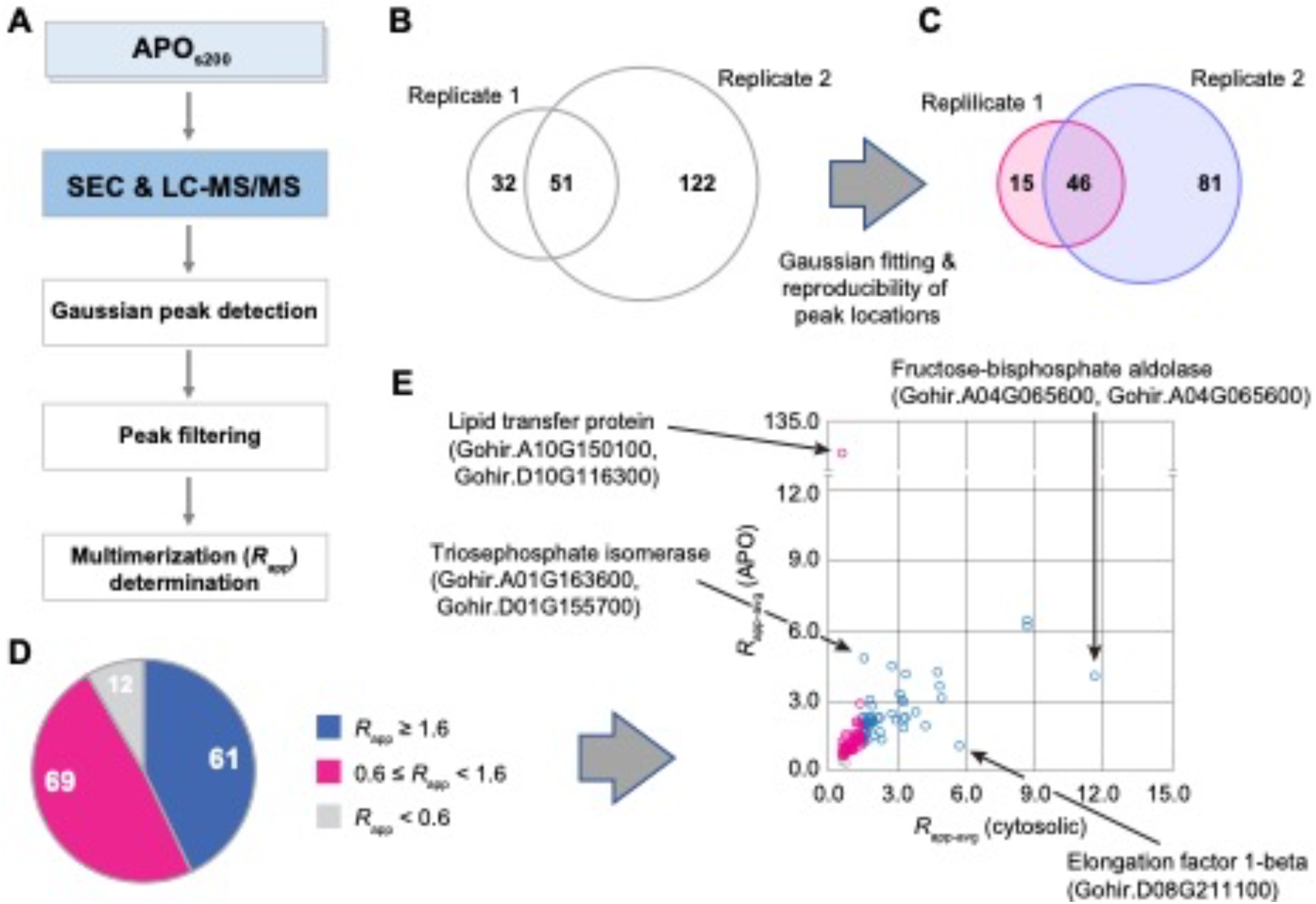
Intactness and multimerization states of dual localized apoplastic proteins. **A,** The CFMS pipeline combined with size exclusion chromatography (SEC) is used to analyze the intactness and potential multimerization states of soluble apoplastic (APO_s200_) proteins. **B,** Coverage of proteins identified in chromatography fractions from two replicate separations. **C,** High confidence peaks present based on 2-fraction shift between the two replicates. **D,** APO_s200_ proteins are intact and are frequently predicted to be in a multimeric state. *R_app_* is used as a diagnostic for multimerization. **E,** Comparison of the apparent mass of 104 apoplast-localized proteins (APO_s200_) with those that had previously reported *M*_app_ measurements from the S200 fraction (Lee and Szymanski, 2021). A/D suffixes reflect ambiguity with respect to homoeolog identification and D indicates homoeolog-specific peptides were identified.

104 APO_s200-SEC_ proteins also had reported *M*_app_ in a previously published SEC profiling experiment of cytosol-enriched proteins isolated from cotton fibers at the same developmental stage (Lee and Szymanski, 2021). Most of the measured *M*_app_ values for APO_s200-SEC_ matched the cytosol-enriched pool and fell along the diagonal (Figure 3E). However, there were several instances in which the multimerization state of a protein differed between the APO_s200-SEC_ and cytosol enriched fractions. For example, a lipid transfer protein (Gohir.A10G150100 and Gohir.D10G116300.) and a triose phosphate isomerase (Gohir.A01G163600 and Gohir.D01G155700) had higher assembly states in the APO_s200-SEC_ fraction. The converse was true for the translation factor EF-1B (Gohir.D08G211100) and aldolase (Gohir.A04G065600 and Gohir.A04G065600). Perhaps these reflect distinct functions in these two compartments.

### Analyses of the soluble particulate and proteomes in the apoplastic fraction

To further characterize the apoplastic proteome, we scaled up the isolation to 4 locules and subjected APO_T_ to ultracentrifugation to generate particulate (APO_p200_) and soluble (APO_s200_) fractions (Figure 4A). There was a visible pellet in APO_p200_ and its protein was ∼63 μg/boll, 4 times higher compared to APO_s200_. To test for the presence of apoplastic vesicles, the APO_p200_ particles were resuspended and processed for negative staining and transmission electron microscopy using contrast agents that interact with biological membranes (Brenner and Horne, 1959). The APO_p200_ contained numerous spherical particles that ranged in size from ∼60 to ∼350 nm (Figure 4B). Similar size ranges for extracellular vesicles have been previously reported (Regente et al., 2017; Rutter and Innes, 2017). These results indicated that the APO_p200_ proteome was dominated by lipid-enclosed particles that may correspond to extracellular vesicles. We will refer to them simply as apoplastic vesicles in this paper.

**Figure 4.**
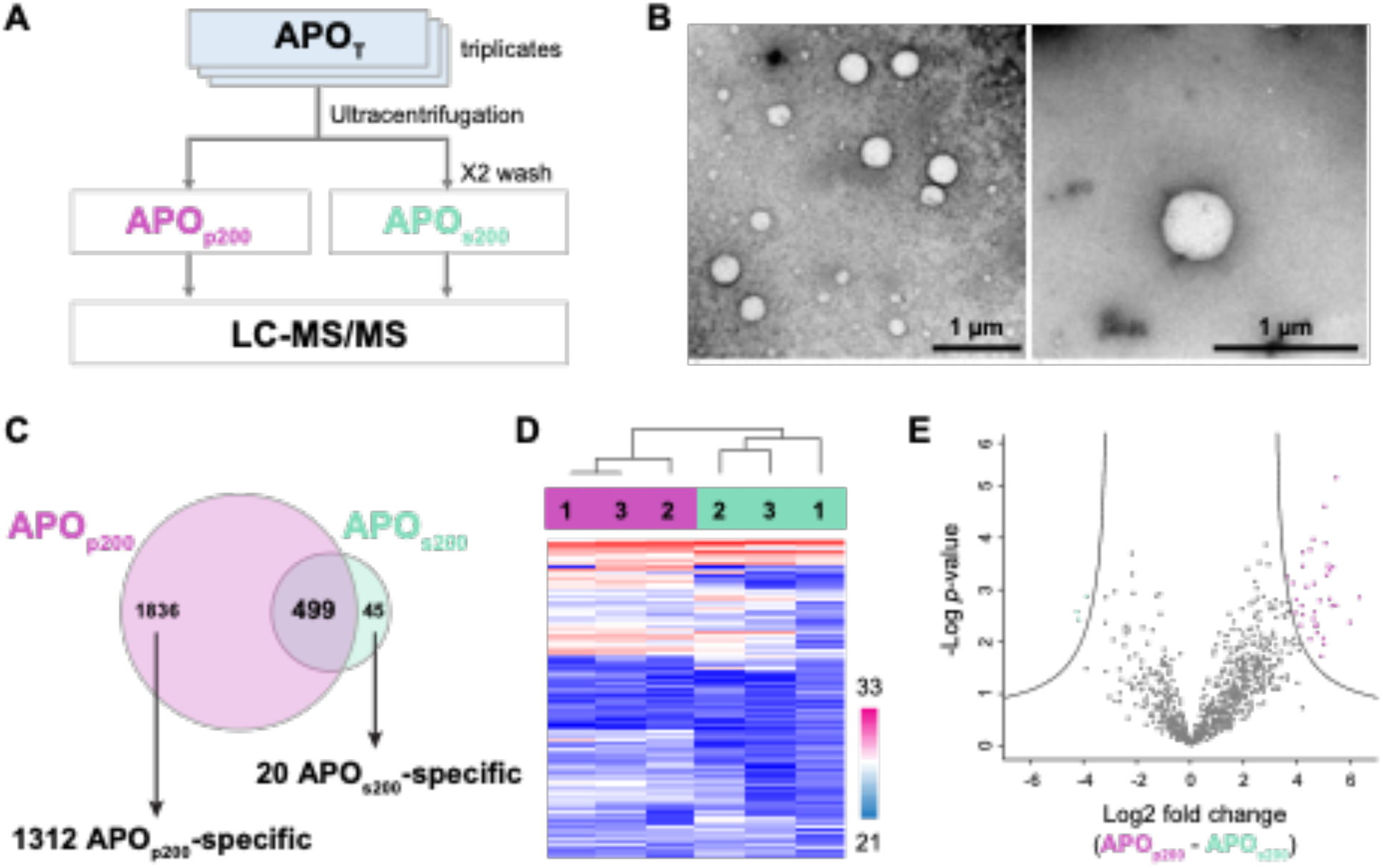
Separation and identification of soluble (APO_s200_) and vesicle-associated (APO_p200_) fractions in the apoplast (APO_TOTAL_). **A,** The pipeline used to identify APO_p200_ and APO_s200_ proteins. **B,** TEM images of vesicles present in the apoplast cellular fraction. **C,** Number of proteins reproducibly identified by MS/MS counts in the APO_p200_ and APO_s200_ fractions. Fraction specific proteins were identified based on MS/MS count data across the replicates and fractions. **D,** Hierarchical clustering and heatmap showing relative protein expression (log2-transformed LFQ protein intensities) of the proteins detected in both apoplast fractions. Columns correspond to the three replicates of both fractions. Missing values were imputed by normal distribution (width = 0.3, shift = 1.8) using Perseus version 2.0.7.0. **E,** Volcano plot depicting two-sample *t*-test result between all quantified proteins present in D. The lines indicate 1 % FDR and 8-fold changes.

Triplicate APO_p200_ and APO_s200_ samples were analyzed using LC/MS, identifying an additional 2335 apoplast localized proteins that were not identified previously (Supplemental Datasets S9 and S10). There was compositional overlap between the two fractions, but APO_p200_ had a much more diverse proteome, having 1312 APO_p200-specific_ proteins based on the spectral count criteria used above (Figure 4C and 4D). It is the APO_p200_ fraction that is driving the similarity between APO_T_ and p200. The APO_s200_ fraction was dominated 499 dual localized proteins. These dual-localized proteins are likely to be peripherally associated with microsomes and only 50 were predicted to be integral membrane proteins based on bioinformatic screens for membrane spanning segments using DeepTMHMM (Hallgren et al., 2022). In the analysis, 267/499 were predicted to be globular and cytosolic (Supplemental Dataset 10A). In the APO_s200_ fraction, 392 proteins were quantified in at least 2 out of 3 replicates, and 96 of them were also quantified in the s200 fraction. Abundances of these dually localized proteins were varied in both fractions, while the abundance ranges were similar, suggesting APO_s200_ protein yield was similar to s200 (Supplemental Dataset S10E). The LFQ intensity-based comparisons of dual-localized proteins identified 35 and 3 upregulated proteins in the APO_p200_ and APO_s200_, respectively (Figure 4E; Supplemental dataset S10C). Another two LTPs and a secreted glycosyl hydrolase were identified as APO_S200-enriched_, and the 35 APO_p200-enriched_ proteins included a diverse set of enzymes, transcription factors, and vesicle coat proteins (Supplemental Dataset S10C).

Having data on the APO_s200_ and APO_p200_ fractions allowed us to compare the predicted organelle localization among all of the cell fractions generated in this study (Table 1; Supplemental Dataset S10A). We limited the analysis to the subset of proteins in each cell fraction that had a unique SUBAcon localization prediction to a specific organelle and calculated the proportion of proteins residing in each cellular locale. As expected, no clear differences were apparent among APO_T_, s200, and p200. Also as expected, the APO_s200-specific_ protein population was highly enriched in predicted extracellular proteins, including a collection of LTPs, cell wall proteins, glycosyl hydrolases, peroxidases, and proteases (Supplemental Dataset 10A). The proteins classified as “enriched” or “specific” in APO_T_ relative to s200 and p200 also had an elevated representation of extracellular proteins compared to other cell fraction datasets. The vacuole was the only compartment that had an increased representation in the cell fractions enriched for apoplastic proteins, including the APO_p200-specific_ fraction (Table 1). Potential reasons for this modest enrichment for vacuolar proteins will be discussed below.

**Table 1.** Accounting of SUBAcon localization predictions for the different cell fractions including only proteins that have a predicted SUBAcon localization to a single subcellular compartment (proportion %).

| cell fraction | Total | Cytosol | ER | Apo. | Golgi | Mito. | Nucleus | Perox. | Plasm. | Plastid | Vacuole |
| --- | --- | --- | --- | --- | --- | --- | --- | --- | --- | --- | --- |
| p200 | 2662 | 0.38 | 0.07 | 0.04 | 0.07 | 0.08 | 0.1 | 0.03 | 0.09 | 0.1 | 0.03 |
| s200 | 175 | 0 | 0 | 0 | 0 | 0 | 0 | 0 | 0 | 1 | 0 |
| APO<sub>T</sub> | 1721 | 0.42 | 0.08 | 0.05 | 0.07 | 0.09 | 0.08 | 0.03 | 0.06 | 0.07 | 0.04 |
| APO<sub>specific</sub> | 42 | 0.1 | 0.02 | 0.21 | 0.07 | 0.19 | 0.14 | 0 | 0.05 | 0.1 | 0.12 |
| APO<sub>enriched</sub> | 49 | 0.14 | 0.08 | 0.33 | 0.04 | 0.1 | 0.08 | 0 | 0 | 0.02 | 0.2 |
| APO<sub>p200-specific</sub> | 1882 | 0.29 | 0.06 | 0.03 | 0.08 | 0.12 | 0.14 | 0.03 | 0.05 | 0.13 | 0.08 |
| APO<sub>s200-specific</sub> | 44 | 0.23 | 0 | 0.52 | 0 | 0.05 | 0.02 | 0.07 | 0 | 0.09 | 0.02 |

Apo.: apoplast; Golgi: Golgi apparatus; Mito.: mitochondria; Perox.: peroxisome; and Plasm.: plasma membrane

## Discussion

This paper describes a simple cell fractionation and quantitative proteomics workflow that provides useful information on the abundance, localization, and multimerization states of the cotton fiber proteome. Although the single locule analysis yielded low coverage of the crude cytosolic (s200) fraction, the crude microsome (p200) and apoplastic (APO_T_) fractions included thousands of fiber proteins. These measured protein abundances spanned 5 orders of magnitude and provide a glimpse into the systems of proteins that enable the cells to reproducibly generate long, thin, spinnable fibers that feed the textile industry. The microsomal fraction contains known proteins that relate to cellulose synthesis and the dynamic organization of the microtubule array, and therefore provides a way to analyze how these and to be discovered proteins operate as part of a cytoskeleton-cellulose synthesis-cell shape control module that is central to fiber morphogenesis. All of these data are available in well-organized supplemental tables that are searchable based on both Gorai/Gohir locus IDs, Phytozome-generated ortholog group numbers, and best-hit Arabidopsis ortholog IDs.

The p200/crude microsome fraction had high protein yields and good coverage in the single locule assay. The most abundant p200 proteins point to specific cellular activities and key homoeologs (or homoeologous pairs when the proteomic data are more ambiguous) that are likely to play a central role in fiber development. For example, the actin and microtubule cytoskeletons are required to pattern fiber growth (Bao *et al*., 2011; Graham and Haigler, 2021; Han et al., 2013; Seagull, 1986; Thyssen et al., 2017; Tiwari and Wilkins, 1995; Wang et al., 2009; Wang et al., 2010a; Yu et al., 2019; Zang *et al*., 2021). TUA2/Gorai.009G224800 (Gohir.A05G214700/Gohir.D05G218000) and TUB8/Gorai.003G126300 (Gohir.A03G047400/Gohir.D03G119800) are among the top 15 expressed proteins in p200 and are predicted to assemble into an abundant heterodimer in microtubules at 9 DPA. Additional orthologs predicted to be involved in microtubule cortical array organization and cellulose synthesis are summarized in Supplemental Dataset S2C and S2F. The top 15 included GhACTIN7/Gorai.003G069800 (Gohir.A11G228966/Gohir.D11G231100), GhACTIN11/Gorai.012G070300 (Gohir.A05G349400/Gohir.D04G062900) orthologs. The GhACTIN11 ortholog predicted to correspond to the dominant Ligon lintless-1 (Li1) mutant of GhACT_LI1 (Thyssen *et al*., 2017) was the fifth most abundant actin isoform in p200. Flux of reduced carbon into cell wall polysaccharides is central to cell wall biogenesis and fiber development. The p200 top 15 included UDP-glucose pyrophosphorylase/Gorai.007G188400 (Gohir.A11G169600/Gohir.D11G175800) and Sucrose synthase/Gorai.009G038000 (Gohir.A05G036000/Gohir.D05G037500), two key enzymes in this pathway (Ahmed et al., 2018; Amor et al., 1995; Coleman et al., 2007; Ruan et al., 2003). The top 15 also included enzymes in methionine biosynthesis, homocysteine S-methyltransferase/Gorai.004G251500 (Gohir.A08G219500/Gohir.D08G236300), and two related to S-adenosyl-methionine-based one-carbon metabolism, adenosylhomocysteinase/Gorai.006G223900 (Gohir.A09G206600/Gohir.D09G200200) and serine hydroxymethyltransferase/Gorai.011G153900 (Gohir.A10G127300/Gohir.D10G138600). Methylation reactions during homogalacturonan biosynthesis (Du et al., 2020) and central carbon metabolism (Hanson and Roje, 2001) may be major sources for carbon utilization.

We focused much of our attention on characterizing the apoplastic wash (APO_T_) fraction because of its importance in cell wall biology and its unexpected protein diversity and abundance. The plant apoplast proteome has been analyzed in numerous previous publications (Supplemental Dataset S4). This compartment frequently contains both soluble proteins and a particulate fraction that can include both membrane enclosed vesicles and large protein/RNA complexes that can sediment during ultracentrifugation (Borniego and Innes, 2023; Zand Karimi et al., 2022). We used ultracentrifugation conditions that do not efficiently sediment large protein complexes like Rubisco (Aryal *et al*., 2014). Negative staining electron microscopy, confocal microscopy, and ultracentrifugation of the apoplast demonstrate that our apoplast fraction contains both a soluble pool and a particulate fraction contains membrane-enclosed vesicles (Figures 1, 4 and 5; Supplemental Dataset S10). Across kingdoms, the extracellular proteome is cloaked in mystery and notoriously difficult to analyze due to variability in methodology and the poorly understood contributions of broken or dead cells (Consortium et al., 2017; Rutter and Innes, 2020; Rutter et al., 2017).

For decades numerous localization and cell fractionation papers have reported the unexplained existence of known cytosolic proteins like calmodulins or 14-3-3/GRF proteins in the extracellular space (Ling and Assmann, 1992; Wen et al., 2007). Our apoplastic datasets are also quantitatively enriched in calmodulin, calmodulin-like, and 14-3-3/GRF proteins (Supplemental data S6 and S10). Serine/threonine-protein kinases are known to phosphorylate RNA-binding proteins of the splicing machinery (Rodriguez Gallo et al., 2022). The kinase ortholog GhSRPK3A/Gorai.006G102900 (Gohir.A09G086500/Gohir.D09G086460) was the highest count protein in 2/3 APO_s200_ replicates (Supplemental Dataset S10B) even though it had no predicted signal sequences or TM. The presence of diverse protein types in the apoplast cannot be explained by contamination from broken cells. For dual localized proteins, the APO_T_ abundance are not similar with that in the s200 fraction (Figure 2D). The APO_s200_ fraction was also highly enriched in known extracellular proteins that harbor conventional signal sequences to enable secretion (Table 1; Supplemental Dataset S10). These included numerous glycosyl hydrolases, e.g. GhGH32A/Gorai.008G216800 (Gohir.A12G199100/Gohir.D12G201800) and xyloglucosyl transferase/Gorai.007G095000 (Gohir.A11G084800/Gohir.D11G088900). Even more convincingly, several soluble apoplastic proteins displayed unique multimerization states compared to their crude cytosolic pool (Figure 3E). For example, GhLTP5A/Gorai.011G128200 (Gohir.A10G150100/Gohir.D10G116300) assembled into a large complex but was predicted to be monomeric in the s200. GhLTP5A is one of several high abundance apoplastic LTP/PR/Bet v 1 proteins, e.g. GhLTP1A/Gorai.011G128100 (Gohir.A10G150100/Gohir.D10G116300), GhLTP1B/Gorai.005G038900 (Gohir.A02G027300/Gohir.D02G035200), GhLTP1C/Gorai.011G128300 (Gohir.A10G150100/Gohir.D10G116400), GhBetV1A/Gorai.012G129000 (Gohir.A04G103900/Gohir.D04G143600), GhBetV1B/Gorai.011G197500 (Gohir.A10G170000/Gohir.D10G176900) that may be involved in extracellular non-vesicular transport of a variety of hydrophobic molecules to the extracellular space (DeBono et al., 2009; Morris et al., 2021; Sterk et al., 1991) and cuticle formation (Kim et al., 2012; Kim et al., 2023; Liu et al., 2014; Thompson et al., 2017; Yatsu et al., 1983). The presence of abundant LTPs in all cell fractions implies this class of protein has a general important role in fiber cells.

The origin and presence of easily extracted membrane-enclosed vesicles in the APO_T_ fraction is even more difficult to explain. Unconventional routes for protein transport to the apoplast have been described in the plant literature (Robinson et al., 2016; Wang et al., 2010b), and membrane-enclosed organelles in the apoplast have been observed in electron micrographs in cryo-preserved samples (Akita et al., 2017; De Bellis et al., 2022). However, genetic experiments in yeast (Oliveira et al., 2010) and Arabidopsis (De Bellis *et al*., 2022) indicate a general importance of anterograde trafficking, but no single trafficking pathway has been found to be required for extracellular vesicle production. Like all other previous studies, we cannot rule out a contribution of broken cells to our apoplast protein samples, but if the identities and abundances of proteins in the APO_T_ and APO_p200_ proteomes reflect a true extracellular pool, the data point to a rather non-specific route of vesicle/organelle fragment delivery to the apoplast.

The slight enrichment of vacuole localized proteins (Table 1) might reflect the enhanced ability of the vacuole to fuse with or donate membranes to the extracellular space (Hatsugai et al., 2009). The size distribution of the vesicles is also impossible to reconcile with passive diffusion of vesicles through the wall. Most vesicles diameters range from ∼100-350 nm, and this exceeds the porosity/void space estimates of the cell wall by more than an order of magnitude (Read and Bacic, 1996; Titel et al., 1997; Zheng et al., 2017). One possible explanation is that the apoplastic vesicles are generated during nano-scale cell wall rupture as bundles of densely packed and strongly adherent cells (Singh et al., 2009) grow at different rates as they elongate and bend to fill the volume of the developing boll capsule. This mechanism would explain the non-specific nature of membrane/organelle export as any organelle that is near the site of rupture would spew into the extracellular space. A localized cell wall rupture could also provide an escape route for the vesicles into to an unrestricted apoplastic space.

This subcellular proteomics identified the pool of apoplast proteins that are not degraded and displays complex multimerization behaviors. Also, the apoplast proteome resembles that of crude microsomes but has enriched protein markers. Those identified APO proteins are present in many anabolic pathways, including RuBisCO shunt and pathways, such as gluconeogenesis, glutamine biosynthesis, pentose phosphate pathway, that involve in NADH and NADPH concentrations (Supplemental Figures S3 and S4). Interestingly, malate dehydrogenase (1.1.1.40 and 1.1.1.37), isocitrate dehydrogenase (1.1.1.42 and 1.1.1.41/1.1.1.286), glucose-6-phosphate dehydrogenase (1.1.1.49), and phosphogluconate dehydrogenase (1.1.1.44) were present in APO_T_ and APO_p200_ fractions, suggesting specific functions within the apoplast. These enzymes provide the reducing power necessary for biosynthetic pathways, such as fatty acid and cell wall biosynthesis, which are essential for fiber elongation. Presumably, the apoplast vesicles could serve as localized hubs for NADPH production, and this localized energy supply could be more efficient for supporting specific cellular activities like fiber elongation, where demand might be high.

### Improvements and future challenges

The proteomic method described here provides a new way to gain improved coverage and crude estimates of sub-cellular localization by analyzing coupling of quantitative proteomics with cell fractionation. There is great potential to use this proteomic based pipeline to analyze fiber development over time, across genotypes, or as a function of environmental conditions. The single locule method here was not sufficient to analyze of the crude cytosolic/s200 fraction.

Either more input material or improved protein precipitation methods will be needed to get acceptable coverage of this important subcellular compartment. The data here suggest that fibers have a complex extracellular proteome that includes more than just proteins that are secreted through standard anterograde trafficking. Based on the intactness and unique multimerization of apoplastic proteins, these are not simply degraded proteins. Further research is needed to characterize the chemistry and potential compartmentalized metabolism of the apoplast and how it might relate to cell wall remodeling and coordinating growth within and between cells.

## Methods

### Cotton cultivars and growth conditions

Cotton plants (*Gossypium hirsutum* cv. TM1) were cultivated in a Conviron^®^ E15 growth chamber (Conviron) at the College of Agriculture Plant Growth Center in the Purdue University. Seeds were sown in 3.0 gallon pots with a soil mixture prepared as 4:2:2:1 soil:perlite:bark:chicken grit. The growth chamber was controlled to generate 50% relative humidity and remain with a day/night setting of 28/23 °C and 16/8 h at a light intensity of 500 μmol m^−2^ s^−1^ (fluorescent lamps of 28 Sylvania F72T12/CW/VHO 100W and 4 Sylvania F24T12/CW/HO 35W; incandescent bulbs of GE 60W light 130V A19). The 30 mins two-step ramp-up and 30 mins two-step ramp-down periods (15 minutes at 166 μmol photons + 15 minutes at 336 μmol photons) were programmed at the daytime beginning and the ending, respectively. Cotton flowers were marked at anthesis as 0-day post anthesis (0 DPA) and harvested at 9 DPA. The harvested cotton bolls were maintained on ice and dissected immediately to obtain intact ovules from each locule.

### Collection of apoplast fluid fraction (APO_T_)

The fibers in isolated locules were fragile and did not tolerate vacuum infiltration and high salt buffers that have been used to isolate extracellular fluids. Therefore, individual locules (∼700 mg each) were dunked in 5 mL of microsome isolation buffer (MIB) [50 mM HEPES/KOH (pH 7.5), 250 mM sorbitol, 50 mM KOAc, 2 mM Mg(OAc)_2_, 1 mM EDTA, 1 mM EGTA, 1 mM dithiothreitol (DTT), 2 mM PMSF and 1% (v/v) protein inhibitor cocktail (160 mg/mL benzamidine-HCl, 100 mg/mL leupeptin, 12 mg/mL phenanthroline, 0.1 mg/mL aprotinin, and 0.1 mg/mL pepstatin A)] and incubated for 10 minutes under gentle shaking. The resulting solution was recovered through 2-layers of cheesecloth and then the entire APO fluid (APO_T_) cellular fraction was used for protein precipitation using an acetone precipitation method. Three biological replicates were prepared.

### Isolation of cytosolic (s200) and microsome (p200) fractions

From the material used for the APO_T_ fraction collection, fibers were directly collected as described previously (Aryal *et al*., 2014; Lee and Szymanski, 2021; McBride *et al*., 2019; McBride *et al*., 2017). Using forceps, fibers were isolated from seeds without any damage to the seeds under 1 mL of MIB solution (sample to MIB ratio was 1:4). The fresh fiber tissues were ground under the cold MIB using Polytron homogenizer (Brinkman Instruments) with chilled blade tip with 10 seconds grinding, 1 minute rest on ice, and another 10 seconds grinding. The homogenate was filtered with 4 layers of cheesecloth pre-soaked in cold MIB, and the cheesecloth was further squeezed to get a residual sample in the cheesecloth. Then the filtered homogenate was further centrifuged on an Allegra X-30R centrifuge (Beckman Coulter Life Sciences) at 1,000 x g for 10 min to remove debris. The supernatant was fractionated at 200 k x g for 20 minutes at 4°C using a Beckman Optima Ultracentrifuge with TLA110 rotor (Beckman Coulter Life Sciences) into a cytosolic (s200) fraction and pellet. The entire s200 fraction was used for protein precipitation using acetone precipitation method. To get a microsome fraction (p200), the resulting pellet was washed with MIB, incubated on ice for 10 minutes, and ultracentrifuged at the same condition as shown above. The supernatant was discarded, and this washing step was repeated. Three biological replicates were prepared for both cellular fractions.

### Determination of protein concentration

For the APO_T_ and s200 cellular fractions, dried pellets from the acetone precipitation were dissolved and denatured in 100 μL of 8 M Urea for 1 hour at room temperature. For the p200 fraction, 200 μL of 8 Urea was directly added over the final pellet and incubated for 1 hour at room temperature to denature proteins from membranes. Undissolved debris was removed by centrifugation at 12 k x g for 15 min using an Allegra X-30R centrifuge (Beckman Coulter Life Sciences). Protein assay was performed using BCA assay kit, according to the manufacturer’s protocol (Thermo Fisher Scientific Inc.). Protein yields for fresh fiber weight were 0.015 mg/g, 0.25 mg/g, 0.012 mg/g in APO_T_, p200, and s200, respectively.

### Separation of soluble (APO_s200_) and particulate (APO_p200_) subfractions of APO_T_ by ultracentrifugation

The APO_T_ fraction was obtained from 4 locules in the same boll as described in the APO_T_ isolation method. Apoplastic extracellular vesicles (APO_p200_) were enriched from the APO_T_ using a Beckman Optima Ultracentrifuge at 200 k x g for 20 minutes at 4°C. The supernatant (APO_s200_) was used as a control in a quantitative proteomics analysis for APO vesicle and cargo protein identification. To deplete weakly associated soluble proteins, the pellet was washed twice with 10 mL of MIB as shown above. The APO_p200_ proteins were solubilized with 150 μL of 8M Urea and then ultracentrifuged to remove insoluble materials from the APO_p200_. Triplicates were prepared.

### Apoplast protein complexes for SEC and Co-fractionation mass spectrometry

For size exclusion chromatography, APO_T_ proteins, that were obtained from four locules from one cotton boll, were pooled and then centrifuged to get APO_s200_ as described in the above APO_p200_ experiment. Size exclusion chromatography (SEC) was done as described previously (Lee and Szymanski, 2021). Briefly, 100 μg of APO_s200_ proteins were resolved in a Superdex Increase 200 10/300 GL (GE Healthcare) on an AKTA FPLC system (GE Healthcare). The mobile phase was [50 mM HEPES-KOH pH 7.5, 100 mM NaCl, 10 mM MgCl_2_, 5% glycerol and 1 mM DTT] and flow rate was set 0.65 ml/minute. Fractions (APO_s200-SEC_) were collected at every 500 mL of elution volume. The column was calibrated using the gel filtration kit 1000 (MWGF1000, Sigma-Aldrich Cor.), and the void fraction was determined using blue dextran. Twenty-seven fractions were collected including the first two void fractions. A cold acetone method was used for precipitating proteins in each fraction. Two biological replicates were prepared.

### LC-MS/MS sample preparation

For LC-MSMS analysis, proteins were digested using trypsin as described previously (McBride *et al*., 2019). We used all protein samples from the APO_T_, APO_p200_, APO_s200_, APO_s200-SEC_, s200 preparations, and 50 μg of proteins in the p200 cell fractions. Denatured protein samples were reduced in 10 mM DTT for 45 min at 60°C, and then alkylated with 20 mM iodoacetamide for 45 min at room temperature under the dark. The urea concentration of the solution was brought down to below 1.5 M by additional ammonium bicarbonate before trypsin (Sigma-Aldrich) digestion. The digestion was proceeded at an enzyme to protein ratio of 1 to 50 at 37 °C. After overnight digestion, trifluoroacetic acid (TFA) was added to end digestion. The digested peptides were purified using C18 Micro Spin Columns (74-4601, Harvard Apparatus). Peptide concentration was measured by BCA assay (Thermo Fisher Scientific Inc.). All the peptide samples were adjusted to equal volume. The most concentrated sample had a peptide concentration of 1 μg/μL, and 1 μL of each sample was injected onto the LC-MS/MS system. For the APO_p200_ and APO_s200_ samples, 1 μg of each sample was injected.

### LC-MS/MS data acquisition

LC-MS/MS analysis was carried out as described previously (Barabas et al., 2019). Peptides samples were analyzed by reverse-phase LC-ESI-MS/MS system using a Dionex UltiMate 3000 RSLC coupled nano System coupled with the Orbitrap Fusion Lumos Tribrid Mass Spectrometer (Thermo Fisher Scientific Inc.). Peptides were loaded onto a trap column (300 μm ID × 5 mm) packed with 5 μm 100 Å PepMap C18 medium, and then separated on a reverse phase column (75 μm ID × 50 cm) packed with 2-μm 100-Å PepMap C18 medium (Thermo Fisher Scientific Inc.). Peptides were resolved at a flow rate of 200 nL/min over a 160-min LC gradient. All the data were acquired in the positive ion mode in the Orbitrap mass analyzer using a higher energy collisional dissociation (HCD) fragmentation scheme. The MS scan range was from 375 to 1,500 m/z at a resolution of 120,000. The automatic gain control target was set as standard, maximum injection time (50 ms), dynamic exclusion 60 s, and intensity threshold 5.0e3. MS data were acquired in data dependent mode with a cycle time of 3 s/scan. MS/MS data were collected at a resolution of 7,500.

### Peptide Identification and Quantification

Andromeda search engine on MaxQuant (version 1.6.14.0) was used for relative protein abundance quantification and protein identification (Cox *et al*., 2014; Tyanova *et al*., 2016a). The search was conducted as described (Lee and Szymanski, 2021). Briefly, the search was conducted with all three cellular fractions obtained from the biological triplicates at a single search, but the three different parameter groups for the three cellular fractions were defined to specify different parameters for each cell fraction. The search parameters were as follows: the match between runs function was set with a maximum matching time window of 0.7 min as default; proteins identified by a single unique peptide were selected; the cotton reference was provided from Dr. Jonathan Wendel; Label-free quantification was selected within each parameter group; all other parameters were set as default. For the APO_p200_ and APO_s200_ search, the same settings were applied. For the APO_s200-SEC_ search, we used the same settings except for the Label-free quantification method.

### Data filtering for protein identification in each cellular fraction (APO_T_, p200, s200)

Two layers of filtering strategies were applied for each cellular fraction dataset to identify cellular fraction specific proteins and proteins enriched in the APO_T_ fraction. First, those proteins identified with MS/MS count(s) ≥ 1 in at least 2 out of 3 replicates were chosen as identified proteins in each cellular fraction (Count-based data filtering; Supplemental Data S2). Here APO-specific (APO_specific_) proteins were determined when any proteins in the APO_T_ dataset had no MS/MS count in both p200 and s200 fractions (Supplemental Dataset S6A). Secondly, quantified proteins with valid LFQ values in at least 2 out of 3 replicates in each cellular fraction were selected from the above count-based datasets. Also, as the benchmark for abundance of proteins in different fractions, the percentile rank of each protein was estimated based on protein abundances in each dataset (Supplemental Dataset S5B). To identify APO-enriched proteins (APO_enriched_), LFQ intensities were log2-transformed and then missing values were replaced using imputation function with as default on Perseus (version 2.0.6.0) (Rudolph and Cox, 2019; Tyanova *et al*., 2016b). Two-sample tests between APO_T_ and p200 or s200 were carried out at 5 % false discovery rate (FDR). The APO_enriched_ proteins were defined as an apoplastic protein that was statistically significant at the tests at 5 % FDR and had 2-fold higher protein abundance than one in both p200 and s200 datasets (Supplemental Dataset S6B).

To identify APO_p200_ proteins, the two-step filtering strategy was also applied. Briefly, the APO_p200_ proteins should be identified at least 2 out of 3 replicates by MS/MS count(s), but not shown in the control. Those proteins present in both the APO_p200_ and APO_s200_ datasets should be statistically significant in a Two-sample *t*-test at 1 % FDR with 8-fold change in abundance. The list of proteins in the APO_p200_ was reported in Supplemental Dataset S10.

### Determination of protein oligomerization states using APO_s200-SEC_

We applied the optimized Gaussian fitting algorithm to convert raw profile data into fitted peaks (McBride *et al*., 2017). The fitted peak locations were used to determine the apparent mass (*M*_app_) values of proteins using the SEC calibration curve created above. If a protein elution profile was not fitted to a Gaussian curve, the global maximum (*G*_max_) was replaced to its peak location. We selected peaks that were present within two-fraction shifts between duplicates and Gaussian fitted peaks that were present in either replicate for further analysis. The protein multimerization state (*R*_app_) was defined as the ratio of the *M*_app_ of a protein to the theoretical monomer mass (*M*_mono_) of the protein (Aryal *et al*., 2014; Lee and Szymanski, 2021; Liu *et al*., 2008). The *R*_app_ of ≥1.6 thresholds indicates that a protein is in a complex. For the *R*_app_ comparison between APO_s200-SEC_ proteins and cytosolic proteins, the previous published cotton fiber dataset was downloaded from Table S2 in Lee and Szymanski (2021), and then the *R*_app_ values were searched using cotton gene IDs.

### Electron and fluorescence microscopy of isolated extracellular vesicles (APO_p200_)

The APO_p200_ pellet isolated from a 9 DPA apoplastic wash solution was isolated as described above and diluted 100 times in phosphate buffered saline (pH 7). One microliter of the resuspended vesicles representing APO_p200_ was negatively stained with 1% uranyl acetate on plasma-treated copper grids. Imaging was performed using an accelerating voltage of 40 to 100kV on a Phillips CM-100 Transmission Electron Microscope. For fluorescence microscopy samples were stained with 1 µL FM4-64 (Thermo Fisher Scientific) for 2 hours at 4 °C, mounted on a slide and visualized using a spinning disk confocal microscope and a 100x Plan-APO 1.46 NA oil-immersion objective. Images were using a spinning disk CSU-X1 confocal head (Yokogawa Electric) and the Prime 95B camera (Photometrix, Tuscon, AZ) mounted on a Zeiss Oberver.Z1 inverted microscope controlled using Slidebook software (Intelligent Imaging Innovation, Denver, CO). FM4-64 was excited with 488 nm light and light was detected using 617/73 bandpass filter. SEM included the use of the Quorum PP3010T cryo system mounted on the Thermo Fisher Apreo 2S scanning electron microscope. The samples were mounted on the cryo holder using a mixture of OCT and aqueous colloidal carbon. Freezing was carried out in nitrogen slush before transferring under vacuum to the preparation chamber attached to the microscope. Sublimation was carried out at −90 °C for 5 minutes followed by sputter coating with platinum. Images were recorded using an accelerating voltage of 5KV and a beam current of 0.1nA.

### Scanning electron microscopy of developing cotton fibers

Locules were collected from a dissected boll at 15 DPA. After dissection, the distal end of the locule was cut with a razor blade rotated roughly 30 degrees off of the proximodistal axis to generate a roughly uniform sample a few millimeters thick while preserving the locule tip. The sample was immediately submerged in liquid nitrogen after attachment to the stub. Samples were imaged on a Thermo Scientific Apreo 2S equipped with a Quorum PP3010T Cryo-SEM Preparation System. Sublimation was performed in 5-minute increments, and additional sublimation steps were used as needed to remove surface ice.

### Apoplast wash following capsule wall dissection and locule dunking

As an alternative method to isolate an APO wash sample, outer capsule wall of a 15 DPA boll was carefully dissected without the underlying fiber cells (Supplemental Figure S1). The dissected boll was rinsed in a beaker of deionized water at room temperature. Then the stem was used to suspend the boll in 25 mL of water so that it did not touch the beaker, the exposed locules were submerged, and the cut edge of the capsule wall was above the waterline. Three 10-minute rinses were done, and then a final 1-hour rinse. 25 mL of each rinse was collected and centrifuged in an ultracentrifuge for 25 minutes at 200,000 x g. The pellets were then each resuspended in 80 µL ddH_2_0. These resuspended particles were stored on ice overnight. A 10 µL aliquot was diluted 2-fold and stained in 1 µM FM4-64 for 2 hours at 4 °C. Then, 1µL of the stained sample was transferred to a slide and mounted with a 25 mm x 25 mm cover glass. To test the possibility that the particles originated from the intact inner capsule wall of the boll, the concave domain of a capsule wall was flooded with 100 µL of buffer or water for 1 hr, taking care to not expose the solution to the cut edge of the capsule wall. Almost no fluorescent particles were detected from this fluid.

### *In vitro* cotton ovule culture

G. hirsutum ovaries at 1 DPA were surface sterilized with 80% ethanol, and 3 or 4 ovules were placed into each well of 6 well plates with 4 mL of modified BT culture media per well (Beasley and Ting, 1973). The ovules were cultivated at 30 °C, and the culture fluid was changed approximately every 5 days. At 22 DPA, 3 mL of culture fluid was withdrawn from each well, carefully pipetting up and down 5 times first. The fluid was centrifuged at 1,000 x g for 10 minutes at 4 C using an Allegra X-30R centrifuge (Beckman Coulter Life Sciences). The resulting supernatant was centrifuged at 200,000 x g for 30 minutes at 4 °C using a Beckman Optima Ultracentrifuge with TLA110 rotor (Beckman Coulter Life Sciences). The pellet was resuspended in 30 µL of cold ddH_2_0 and stored in −80 °C until further use.

Five microliters of aliquots from samples with visible pellets were diluted 2-fold and stained with 1 µM FM4-46 overnight at 4 °C. Then 1 µL (1.4% of the resuspended pellet) was placed on a slide, covered with a cover slip (25 mm x 25 mm) and imaged with a spinning disk confocal microscope. Excitation was with 488 nm light and was detected using a 617/73 bandpass filter.

### Comparison of cotton APO_T_ proteins with prior proteomics studies

We created a combined list from 13 published apoplast/extracellular vesicle proteomics studies to validate the results and obtain a clearer picture of the progress made in the secretome field to date. To generate combined list, 12 external datasets from Arabidopsis and 1 from rice secretome studies were used and duplicates from each dataset were eliminated (Supplemental Data S4). All of the Arabidopsis locus IDs were converted to cotton locus IDs by performing the ortholog mapping as described previously (Lee and Szymanski, 2021). To identify overlaps for each protein and to find cotton specific secretory proteins, the list of 1643 APO_T_ proteins was compared with the combined list generated from external datasets (Supplemental Data S4B). The percent overlaps were calculated by dividing the number of Arabidopsis ID commonly found in APO_T_ dataset and external dataset by the total number of Arabidopsis IDs identified in each external dataset.

### Gene ontology enrichment analysis

Gene Ontology (GO) enrichment analysis was performed using the Singular Enrichment Analysis (http://systemsbiology.cau.edu.cn/agriGOv2/index.php) on agriGO v2.0 (Tian et al., 2017). The enrichment was calculated using Fisher’s exact test against Cotton D locus ID v2.1 (Phytozome v11.0) background. Minimum number of mapping entries was 5, and Yekutieli (FDR under dependency) was used as a multi-test adjustment method. Significantly enriched GO terms in biological process (BP), GO molecular function (MF) and GO cellular component (CC) categories were reported at 5% FDR.

### Statistic tests and data analyses

Statistical analysis was performed using R version 4.2.0 (R Core Team, 2018) on RStudio 2022.07.1 (RStudio Team, 2018). Gaussian fitting (https://github.com/dlchenstat/Gaussian-fitting) was applied using MATLAB_R2022a. Microsoft Excel on Office 365 for Mac was used to organized and display the analyzed data.

## Supporting information

Data S1 raw abundances of peptides and proteins

Data S2 APO P200 S200 proteins

Data S3 GO enrichment analysis of proteins in 3 cellular fractions

Data S4 Proteome coverage of APO and published apoplast datasets

Data S5 APOp200 and APOs200 yield different buffers

Data S6 APO-specific and APO-enriched proteins

Data S7 Raw abundance profiles of peptides and proteins in APOs200 (APO SEC)

Data S8 APOs200 protein complex profiling (APO SEC)

Data S9 Raw abundance profiles of peptides and proteins in APOp200 APOs200

Data S10 APOp200 and APOs200 cell fractions

Data S11 GO enrichment analysis of proteins in APOp200 and APOs200

## Data availability

All LC/MS .raw files are annotated and made available at Pride (accession code: PXD051721).

## Acknowledgements

We would like to thank Chris Gilpin and Robert Seller at the Purdue Life Sciences Microscopy Facility (Facility RRID SCR_022687) for assistance with negative stain TEM and cryoSEM. We thank Uma Aryal at the Purdue Proteomics Facility for running the LC/MS samples. Heena Rani was supported by Indian Science and Engineering Research Board (SERB)-Purdue University Overseas Visiting Doctoral Fellowship.

---

**Supplemental Figure S1.**
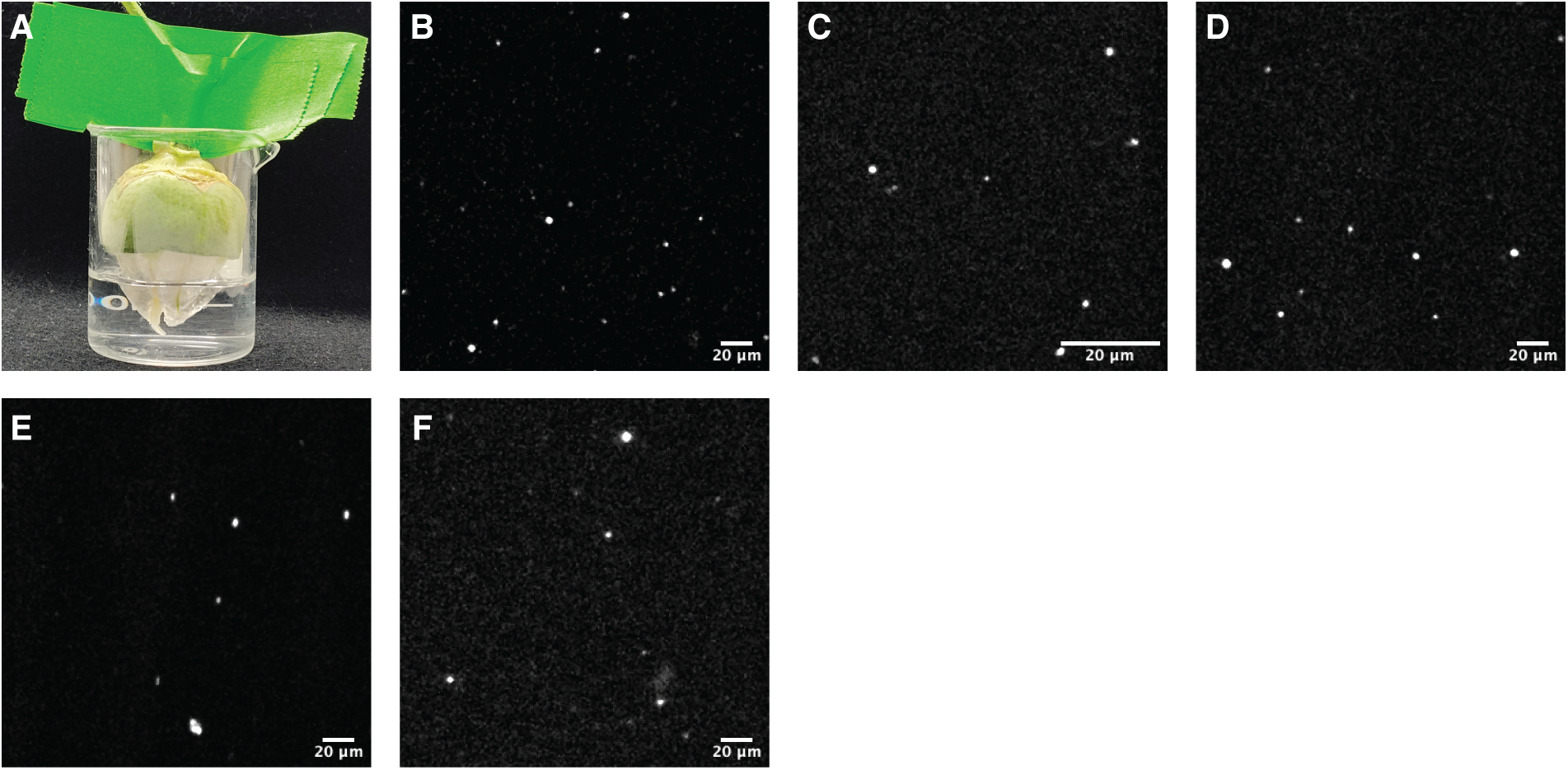
FM4-64 positive vesicles in the apoplast wash following capsule wall dissection and locule dunking. **A,** A set-up for collecting vesicles from the partially dissected 15 DPA boll. **B and C,** FM4-64 positive vesicles after 10 min incubation (1st). **D,** FM4-64 positive vesicles after another 10 min incubation (2nd). **E,** FM4-64 positive vesicles after 10 min incubation (3rd). **F,** FM4-64 positive vesicles after 1 hr incubation (4th).

**Supplemental Figure S2.**
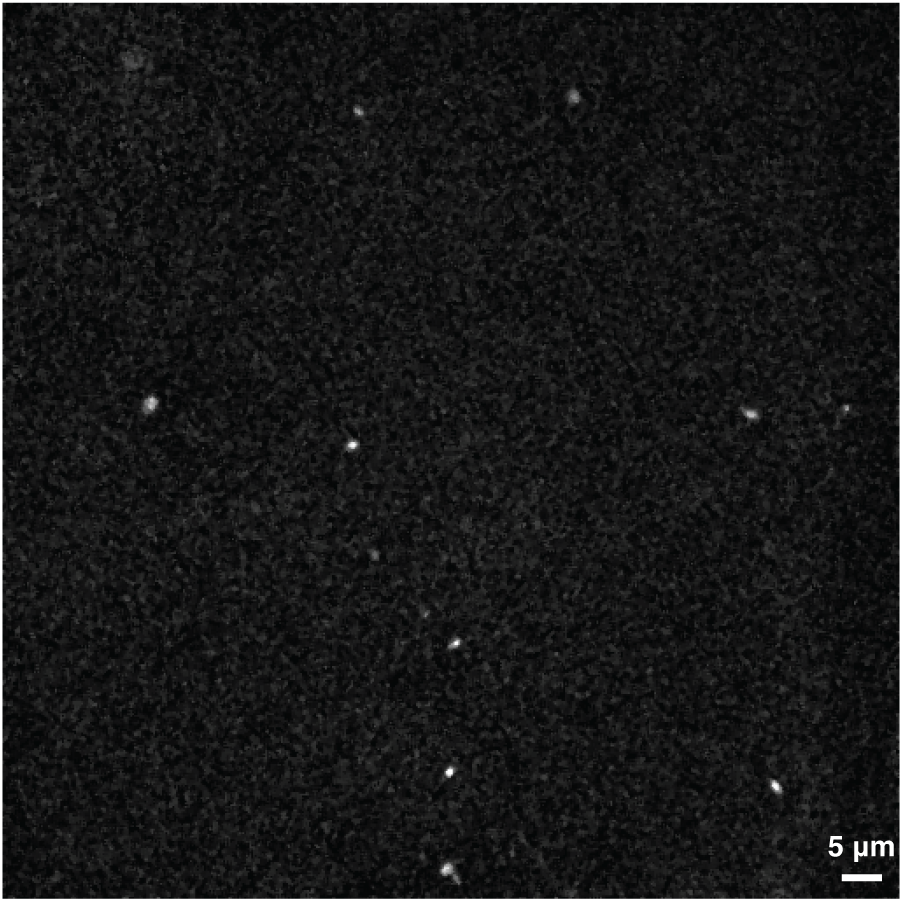
FM4-64 positive vesicles in the culture fluid of cotton ovules at 22 DPA.

**Supplemental Figure S3.**
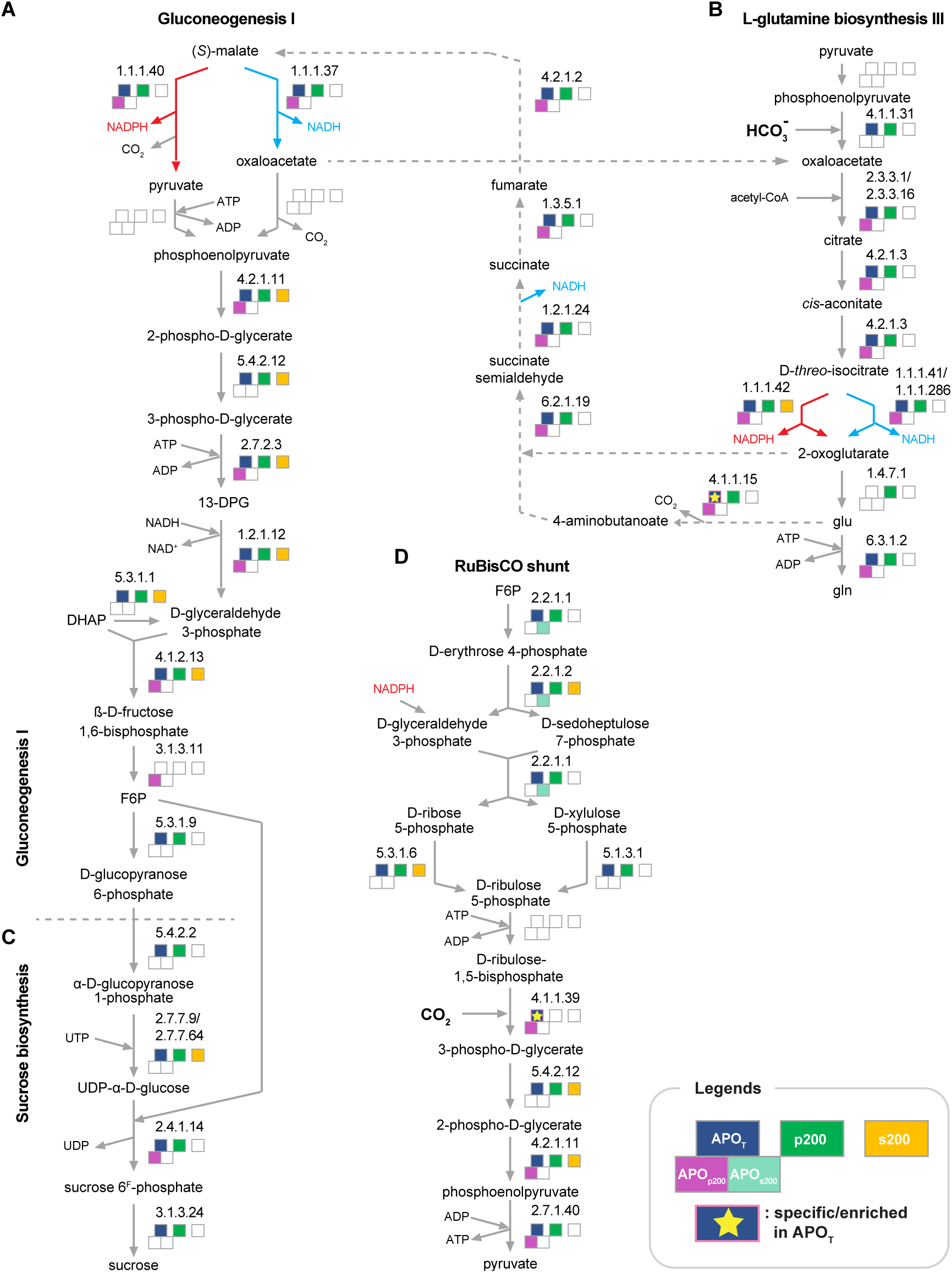
Functional vesicles for catabolic and anabolic pathways in apoplast? Many of proteins in Gluconeogenesis I (A), L-glutamine biosynthesis III (B), Sucrose biosynthesis (C), and RuBisCO shunt (D) were identified in the APO_T_, APO_p200_ and APO_s200_ datasets. Each proteome was projected to metabolic pathways sing Plant Metabolic Network (https://plantcyc.org/).

**Supplemental Figure S4.**
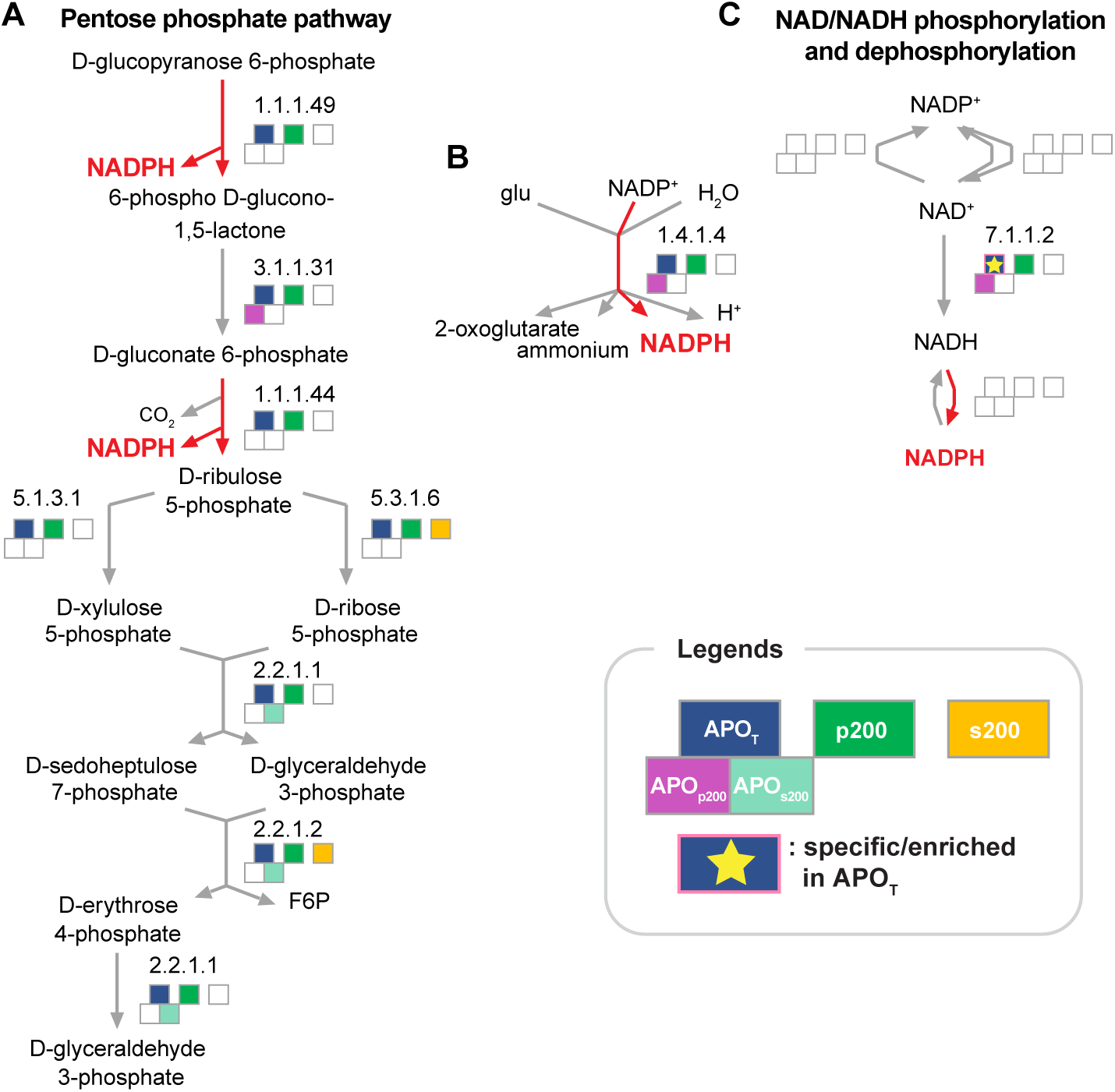
Additional pathways that concentrate NADPH concentrations. **A to C,** Enzymes functioning in NAD and NADPH synthesis are highlighted. Enzymes in each dataset were projected to metabolic pathways sing Plant Metabolic Network (https://plantcyc.org/).

## Supplemental Datasets

**Supplemental Dataset S1.** Raw abundances of peptides and proteins identified in the APO_T_, p200, and s200 of cotton fibers at 9 DPA.

**Supplemental Dataset S2.** Proteins identified and quantified in the APO_T_, p200, and s200 cellular fractions.

**Supplemental Dataset S3.** Gene ontology enrichment analysis of APO_T_, p200, and s200

**Supplemental Dataset S4.** Overlap between APO_TOTAL_ proteome in this study and proteomes of the extracellular space in the previous 13 proteomics studies.

**Supplemental Dataset S5.** APOp200 and APOs200 protein yields in different buffers

**Supplemental Dataset S6.** APO_specific_ and APO_enriched_ proteins.

**Supplemental Dataset S7.** Raw profiles of peptides and proteins identified in the APO_s200_ cellular fraction of the 9 DPA cotton fiber by CFMS (APO_SEC_).

**Supplemental Dataset S8.** Apoplastic protein complex profiling analysis by CFMS (APO_s200-SEC_).

**Supplemental Dataset S9.** Raw abundances of peptides and proteins identified in the extracellular vesicle (APO_p200_) samples.

**Supplemental Dataset S10.** Proteins specifically identified in APO_p200_ and APO_s200_.

**Supplemental Dataset S11.** Gene ontology enrichment analysis of APO_p200_, APO_s200_.

## Abbreviations

DPA: day post anthesis

APO_T_: apoplast wash/fluid fraction

p200: membrane-associated fraction

s200: cytosolic fraction

APO_specific_: apoplast-specific

APO_enriched_: apoplast-enriched

APO_p200_: apoplast membrane-associated fraction

APO_s200_: apoplast soluble fraction

APO_T_-vesicles: membrane-enclosed particles

APO_s200-SEC_: size exclusion chromatography fractions of APO_s200_

*R*_app_: protein multimerization state

*M*_app_: apparent mass of protein

*M*_mono_: theoretical mass of protein

